# Relationships of scale cortisol content suggests stress resilience in freshwater fish vulnerable to catch-and-release angling in recreational fisheries

**DOI:** 10.1101/2022.11.15.516354

**Authors:** J.R. Britton, D Andreou, M Lopez-Bejar, A Carbajal

## Abstract

The capture by angling of an individual fish is recognised as a short-term physiologically stressor. In fish populations exploited by catch and release angling (C&R), there is potential for some individual fishes to be captured on multiple occasions, but the longer term physiological consequences of this remain uncertain. Using scale cortisol content as a biomarker of chronic stress and scale samples from two fish populations exploited by C&R angling, we developed proxies of angling capture vulnerability before testing these proxies against scale cortisol content. In a riverine population of European barbel *Barbus barbus*, fish with the highest scale cortisol content were predicted as those sampled by angling rather than electric fishing, as angled fish had significantly smaller home ranges and diets based primarily on angling baits. In a population of common carp *Cyprinus carpio* in a small pond fishery, we predicted that fish with the highest scale cortisol content would be those with higher proportions of angling bait in their diet. In both species, however, the fish predicted to be most vulnerable to angling capture had the lowest levels of scale cortisol content. We suggest that this is through fish that are captured regularly being highly stress resilient (with this independent of other traits) or fish with traits that suggest high capture vulnerability being able to minimise their recapture rates through developing hook avoidance behaviours after an initial capture. Overall, these results suggest that scale cortisol content is a useful biomarker for measuring chronic stress from C&R angling.

**Highlights:**

- In catch-and-release angling (C&R), a single capture event is a short-term stressor
- The effect of multiple capture events on individual fishes in C&R is unknown
- Scale cortisol content was used as a biomarker of chronic stress
- Fish of high vulnerability to capture had relatively low scale cortisol content
- Fish with high exposure to C&R appear to have high stress resilience.

## 1. Introduction

Global estimates of the number of recreational anglers vary, but suggestions are of up to 700 million recreational anglers in the world who capture 12% of the global fish harvest, primarily from freshwaters and inshore areas (Arlinghaus and Cooke, 2009). Harvesting by angling can result in exploited populations comprising of low-activity, highly stress-responsive phenotypes due to be ‘angling induced selection’ for specific traits, with the most vulnerable individuals to capture often being active phenotypes of high stress resistance (Koeck et al., 2019; Monk et al., 2021). Catch and release angling (C&R) is increasingly being adopted in many world regions as it minimises angling impacts on target populations as captured fish are returned alive, so maintaining phenotypic diversity and providing conservation benefits in exploited populations (Arlinghaus et al., 2007).

In populations exploited by C&R, the capture vulnerability of individuals is also non-random, with some individuals rarely being captured, but with others re-captured on a regular basis (Lennox et al., 2017). For example, across 46 tagged European catfish *Silurus glanis* in a pond fishery, 30 individuals went uncaptured across an entire year, but 8 individuals were captured between 10 and 26 times, with some being recaptured on successive days (Britton et al., 2007). Differences in the capture vulnerability of individuals is associated with intra-population variability in behavioural traits, although the phenotypes most vulnerable to C&R can be species-specific (Alos et al., 2012; Lennox et al., 2017) and, in some species, behaviours such as boldness are not a good predictor of capture vulnerability (Vainikka et al., 2016).

The capture by angling of an individual fish is considered as a short-term physiologically stressful event, with the probability of sub-lethal effects occurring being influenced by a range of individual and interacting variables, including fish size, hook damage extent, fight time, air exposure, general fish handling and environmental conditions (Muoneke and Childress, 1994; Cooke and Suski, 2004; Pinder et al., 2019). Stress responses to single capture events have been detected through changes to blood chemistry (e.g. Cooke et al., 2013) and the application of simple reflex action impairment indicators as a ‘whole-body’ stress response (e.g. Pinder et al., 2019). However, the longer-term physiological effects (i.e. chronic stress effects) of an individual fish being repeatedly captured by C&R angling remain unclear, especially in large, open systems where knowledge on previous capture events of an individual fish might be unknown. In these situations, proxies of individual vulnerability to angling capture could be used to infer capture history. For example, it could be assumed capture vulnerability will be increased in individual fish that have diets more heavily subsidised by angling baits (increasing their probability of encountering a baited hook) (Britton et al., 2022) and/or that have smaller home ranges (increasing their spatial encounters with anglers) (Gutmann Roberts et al., 2019).

The level of cortisol in an individual fish is increased when they are exposed a stressor (Carbajal et al., 2018, 2019a,b). Circulating cortisol levels in fish correlate strongly with scale cortisol content, but with rates of accumulation and clearance in scales being much slower than for blood plasma (Laberge et al., 2019). Although exposing individual goldfish *Carassius auratus* to a single acute air emersion stressor did not influence their scale cortisol content, high and sustained circulating cortisol levels produced from unpredictable chronic stressors did increase it, with this content being heterogenous in scales across the whole body surface (Laberge et al., 2019). Carbajal et al. (2018, 2019a,b) revealed that scale cortisol levels are influenced by mid-term, energetically intense periods rather than long-term stressors, thus can be considered as a biomarker that provides retrospective hormonal measurements from fishes and time periods that are usually difficult or impossible to obtain. Scales have recently been applied to provide a retrospective measure of past stress experience in fishes as diverse as rainbow trout *Oncorhynchus mykiss* (Carbajal et al., 2019a), goldfish *Carassius auratus* (Carbajal et al., 2018; Laberge et al., 2019), Catalan chub *Squalius laietanus* (Carbajal et al., 2019b), sea bass *Dicentrarchus labrax* (Lebigre et al., 2022), and dab *Limanda limanda* (Vercauteren et al., 2022), where the focus has been on responses to aquaculture and environmental pollution. Individual fishes with relatively lower scale cortisol content would thus be assumed to have higher resilience to chronic stress than those with relatively high content.

The aim here was to apply scale cortisol levels as a biomarker of chronic stress to fish exposed to C&R angling in recreational fisheries. This biomarker was applied to two fish populations exploited by C&R angling, European barbel *Barbus barbus* (‘barbel’) in a riverine fishery and common carp *Cyprinus carpio* (‘carp’) in a pond fishery. As the previous capture history of these fishes were unavailable, the initial objective was to develop proxies of vulnerability to angling capture. For barbel, these proxies were based on the sampling method, dietary reliance on angling bait (from stable isotope analysis, SIA), and home range size (from acoustic telemetry). For carp, the proxies were based only on their dietary reliance on angling bait (from SIA). The second objective was to then test these proxies against the corresponding scale cortisol levels. We posit that fish with proxies that suggest high vulnerability to angling (re)capture will have significantly higher scale cortisol content (due to repeated capture events) than fish of low vulnerability.

### 2. Materials and Methods

#### 2.1 Study species

Barbel is a rheophilic species encountered in many European rivers and is a popular angling target species due to their aggregative behaviours that facilitates the capture of multiple individual fish during a single angling event, where fish might be captured between 1 and 8 kg (Britton and Pegg, 2011). Their populations are mainly comprised of individuals that have relatively small home ranges (< 1 km), although a small proportion of individuals are usually more mobile, with home ranges exceeding 10 km ((Britton and Pegg, 2011). In the last 20 years in the study river (see below), barbel angling tactics have moved almost entirely to using manufactured baits, often with a strong marine fishmeal base, where relatively large amounts are released into the water by anglers which are then consumed by the fish (Gutmann Roberts et al., 2017; De Santis et al., 2019). These baits, and those also based on plants (such as maize) tend to be highly enriched in the δ^13^C stable isotope (> -22.0 ‰), whereas the putative prey resources of these fishes tend to highly depleted (< -28.0 ‰), thus providing the ability to assess the extent to which fish are consuming these different resources (*cf*. Britton et al., 2022). Accordingly, SIA has indicated that the diet of some larger barbel (> 400 mm) is primarily comprised of angling baits, whilst other fish of similar size have diets that remain based mainly on natural prey resources (Gutmann Roberts et al., 2017; De Santis et al., 2019).

Introductions of carp for both aquaculture and angling have resulted in their domesticated strains being invasive in many of the world’s freshwaters (Vilizzi et al., 2015), with their popularity for angling resulting from their relatively large size and fighting qualities (Britton 2022). Lentic recreational fisheries tend to be stocked either with very high abundances of smaller fish (e.g. 1 to 7 kg) (North 2002) or smaller numbers of relatively large fish (>10 kg) (Žák 2021). In the study area (see below), angling tactics in these fisheries also tend to be dominated by use of manufactured baits, many of which are maize based with highly enriched values of δ^13^C when compared with natural prey items (Britton et al., 2022; Imbert et al., 2022). Where these fisheries have a high stock density of carp that are fished for regularly, their diet (and the diets of other species present) tend to be dominated by these baits (Britton et al., 2022).

### 2.2 Study waters

Barbel were used here from the lower River Severn basin that were sampled between 2015 and 2020. The scale samples collected in September 2015 were from barbel used in an acoustic telemetry study in which fish were sampled by both electric fishing and C&R angling; the angled fish had significantly smaller home ranges than those electric fished (*cf*. Gutmann Roberts et al., 2019). Barbel scale samples were then also collected in summer 2016, 2018 and 2020, although the home range sizes were not measured for these fish. The full methodology of fish capture, tagging and scale removal are outlined in Gutmann Roberts et al. (2019). In summary, the data recorded for each individual fish was its sampling method (as angling or electric fishing), fork length (nearest mm), and up to 5 scales were taken from the area below the dorsal fin but above the lateral line. All fish were the returned alive to the river. The methodology for relating to the acoustic tagging and estimation of individual home range sizes is available in Gutmann Roberts et al. (2019). At the same time as the fish sampling, amphipod samples (gammarids) were collected as a putative fish prey resource.

Carp were sampled in summer 2020 from a small (1.5 ha) pond in Southern England that is managed for intensive recreational angling. Multiple anglers are present at this fishery on a daily basis for much of the year (e.g. > 20 anglers per day), including participating in competitions that generally last for 5 hours and where captured fishes are held in keep-nets until the end of the competition before being batch-weighed and then returned alive to the water. This fishery had been stocked with fish to provide a very high fish biomass to support angler catches that reach over 10 kg h^-1^ (North 2002), and where relatively high volumes of angling baits are released to enable these catch rates to be maintained during the entirety of the angling session (Britton et al., 2022). The scale samples used here were all from fish captured by angling; the fish were identified to species, measured (fork length, nearest mm), and up to three scales removed. The carp used in analyses were between 530 and 700 mm fork length. After sampling, the fish were then returned alive to the pond. Concomitantly, amphipod samples were also collected as a putative fish prey resource.

All of the fish sampling was completed following ethical review and under UK Home Office licence P47216841 and 70/8063 (under the UK Animals (Scientific Procedures) Act 1986 and associated guidelines).

### 2.3 Stable isotope analysis

Some of the proxies of angling vulnerability were developed from stable isotope analysis (SIA) of scales of the sampled fishes (*cf*. Results). This involved the removal of scale material from the outer edge of the scale (i.e., material produced in recent months; Hutchinson and Trueman 2006), which was then dried to constant mass at 60 ^°^C. The amphipod samples were also dried in this manner. All samples were then analysed at the Cornell University Stable Isotope Laboratory (New York, USA) for δ^13^C and δ^15^N in a Thermo Delta V isotope ratio mass spectrometer (Thermo Scientific, USA) interfaced to a NC2500 elemental analyser (CE Elantach Inc., USA). Analytical precision of the δ^13^C and δ^15^N sample runs was estimated against an internal standard sample of animal (deer) material every 10 samples, with the overall standard deviation estimated at 0.08 and 0.04 ‰ respectively. Ratios of C:N indicated no requirement for lipid normalisation (Winter and Britton, 2021).

As differences in mean vales of δ^13^C and δ^15^N of the amphipod samples from the River Severn were minor across the sampling years (< 1 ‰) then the barbel SI data were used without correction. For carp, to indicate the extent to which individuals had diets based on angler baits, then their δ^13^C data were converted to corrected carbon (δ^13^C_corr_) (Olsson *et al*., 2009):

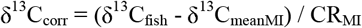

wherein δ^13^C_fish_ is the δ^13^C value of each fish, δ^13^C_meanMI_ is the mean δ^13^C of the macroinvertebrate prey and CR_MI_ is the carbon range (δ^13^C_max_ - δ^13^C_min_) of the same macroinvertebrates (Olsson *et al*., 2009). As discrimination factors of δ^13^C between prey and fish predators are generally 1 to 2 ‰, but can be higher for scale (e.g. up to 5 ‰ on invertebrate based diets; Busst and Britton 2016), then fish values of δ^13^C_corr_ outside of these ranges (e.g. > 5 ‰) would suggest the fish were feeding on alternative dietary items to these prey resources (i.e. angling baits), where the higher the value of δ^13^C_corr_, the greater the dietary reliance on angling bait (*cf*. Britton et al., 2022). To complement δ^13^C_corr_, carp δ^15^N data were converted to trophic position (TP) (Olsson *et al*., 2009):

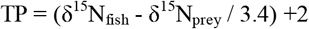

where TP and δ^15^N_fish_ are the trophic positions and the nitrogen ratios of each individual fish, δ^15^N_prey_ is the mean nitrogen ratio of the putative macroinvertebrate prey resource, 2 is the trophic position of these prey resources (as primary consumers) and 3.4 is the generally accepted fractionation factor between adjacent trophic levels (Post, 2002). If the fish had been foraging on the putative macroinvertebrate prey groups used in the TP equation then the fish TP values would be expected to be between 2.5 and 4.5 (with variation resulting from differences in dietary proportions between individual fish) (Busst and Britton, 2016; Winter *et al*., 20121). Values outside of this range would indicate the consumption of alternative dietary items.

### 2.4 Scale cortisol content

Scale cortisol content was determined from scales by enzyme immunoassay (Cortisol EIA KIT; Neogen® Corporation, Ayr, UK). Although the scale mass required to provide reliable cortisol content estimates from this method has not been determined specifically, in other taxa, material from feathers and faeces has required a minimum of =/> 20 mg of material (e.g. Millspaugh and Washburn, 2004; Lattin et al., 2011). For carp, their relatively large scale size and mass meant that the method could be successfully applied on individual fish (scale mass range: 38.7 - 50.9 mg). n = 10). For barbel (n = 68), their smaller scales meant the scale mass available for individual fish was not sufficient for the analysis and so fish had to be pooled in order to provide enough material. To provide groups of fish that were considered to be consistent in their traits and behaviours, they were pooled according to their sampling method (electric fishing or angling), then sampling year (2015/ 2016/ 2018/ 2020) and then by δ^13^C values (< -23.00/ -22.99 to -21.0/ -20.99 to -19.00), resulting in 12 groups of barbel comprising between 3 and 8 fish that provided 20.4 and 50.7 mg of material for analysis (Table 1).

**Table 1.** Meta-data of the groups of European barbel that were pooled for the purposes of scale cortisol content analyses (EF = electric fishing)

| Group | Year | Method | n | Mean length (range)<br>(mm) | Mean $\delta^{13}\text{C}$ (range) (‰) | Mean $\delta^{15}\text{N}$ (range) (‰) | Scale mass<br>(mg) | Scale cortisol<br>content (pg mg<sup>-1</sup>) |
| --- | --- | --- | --- | --- | --- | --- | --- | --- |
| **1** | 2015 | Angling | 6 | 604 ± 47 (557 – 721) | -20.52 ± 0.28 (-20.98 - -20.10) | 11.50 ± 0.24 (11.03 - 11.90) | 45.5 | 2.71 |
| **2** | 2015 | Angling | 8 | 600 ± 41 (529 – 680) | -22.04 ± 0.46 (-22.79 - -21.02) | 12.11 ± 0.59 (10.74 - 13.30) | 50.7 | 2.49 |
| **3** | 2015 | Angling | 5 | 633 ± 40 (585 – 698) | -24.47 ± 1.18(-26.66 - -23.37) | 12.37 ± 0.52 (11.81 - 13.27) | 46 | 1.60 |
| **4** | 2015 | EF | 8 | 584 ± 75 (394 – 717) | -22.08 ± 0.43 (-22.82 - -20.98) | 12.03 ± 0.57 (10.71 - 13.37) | 50.7 | 4.32 |
| **5** | 2015 | EF | 7 | 534 ± 95 (397 – 770) | -24.85 ± 0.91 (-26.50 - -23.25) | 13.27 ± 0.78 (11.81 - 14.88) | 46.3 | 3.73 |
| **6** | 2016 | Angling | 5 | 724 ± 88 (600 – 800) | -20.03 ± 0.50 (-20.55 - -19.37) | 11.33 ± 0.67 (10.48 - 12.31) | 50.2 | 1.64 |
| **7** | 2016 | Angling | 4 | 600 ± 74 (520 – 690) | -22.50 ± 0.72 (-22.95 - -21.40) | 11.85 ± 0.34 (11.47 - 12.25) | 47.9 | 1.71 |
| **8** | 2018 | EF | 3 | 642 ± 86 (576 – 725) | -25.03 ± 0.25 (-25.27 - -24.84) | 13.37 ± 0.45 (13.01 - 13.79) | 38.2 | 5.89 |
| **9** | 2018 | EF | 4 | 537 ± 14 (520 – 550) | -23.95 ± 0.47 (-24.36 - -23.26) | 12.65 ± 0.75 (11.70 - 13.36) | 28.2 | 2.63 |
| **10** | 2020 | Angling | 3 | 618 ± 119 (555 – 739) | -21.53 ± 1.00 (-22.53 - -20.86) | 11.47 ± 0.66 (10.81 - 11.89) | 20.4 | 3.43 |
| **11** | 2020 | Angling | 5 | 707 ± 47 (557 – 721) | -24.64 ± 0.89 (-25.54 - -23.12) | 12.87 ± 0.74 (11.86 - 13.72) | 50.4 | 3.47 |
| **12** | 2020 | Angling | 5 | 626 ± 41 (529 – 680) | -25.45 ± 0.51 (-26.12 - -24.78) | 13.77 ± 0.22 (13.42 - 14.03) | 50.4 | 4.42 |

Cortisol extraction from fish scales was performed following the methodology of Carbajal et al. (2018). In brief, scales were first washed with isopropanol three times to remove any hormone sources that could have been on the external surface of the scales. Once dry, the scale samples of each fish/ group of fish were mechanically pulverized with a ball mill (Retsch, MM2 type, Germany), and each powdered sample was incubated overnight in methanol. After extraction, samples were centrifuged, and the supernatant was evaporated. Dried extracts were reconstituted with enzyme immunoassay buffer provided by the assay kit and immediately stored at −20 °C until analysis.

### 2.5 Data analyses

To develop the proxies for angling vulnerability of barbel, data from the fish sampled in 2015 were used (Gutmann Roberts et al., 2019). The relationships between the sampling method, home range size and stable isotope data of individuals were tested using generalised linear models (GLM), where the dependent variable was home range size, δ^15^N or δ^13^C, the independent variable was sampling method (angling or electric fishing) and fish length was used as the covariate. For carp, the proxies were developed from their relationships between δ^13^C_corr_ and TP according to linear regression.

For barbel, testing differences in scale cortisol content between the groups was then based on the sampling method (*cf*. Results; Table 1) in a GLM, where the dependent variable was scale cortisol content, the independent variable was sampling method, and the covariates were year, mean δ^15^N, mean δ^13^C and mean fork length. For carp, the data could be used continuously, and so cortisol levels were tested against of δ^13^C_corr_ and TP using linear regression.

## 3. Results

### 3.1 Vulnerability to angling capture

In European barbel, the 2015 acoustic telemetry study revealed that the mean (± 95 % CI) home range of fish sampled by angling was significantly smaller than those sampled by electric fishing (2750 ± 1230 m vs 6110 ± 2080 m; GLM: Wald χ^2^ = 6.98, P < 0.01), with length not being a significant covariate (P = 0.15) (Gutmann Roberts et al., 2019). Angler caught fish had significantly enriched δ^13^C values versus the fish sampled by electric fishing (−21.32 ± 0.67 vs -22.99 ± 0.68 ‰; GLM: Wald χ^2^ = 11.89, P < 0.01), with length not being a significant covariate (P = 0.58). Angler caught fish were also significantly enriched in δ^15^N versus those captured by electric fishing (11.77 ± 0.40 vs 12.33 ± 0.40 ‰; GLM: Wald chi-square = 3.84, P = 0.05), with length not being a significant covariate (P = 0.68). Correspondingly, we posit that across all of the samples, groups of fish sampled by angling will have higher vulnerability to angling capture (due to smaller home ranges and so higher spatial encounters with anglers, and their higher dietary contributions of angling baits) and, accordingly, have significantly higher scale cortisol content (due to multiple capture events in preceding months).

In carp, values of δ^13^C_corr_ ranged between 6.22 and 8.10 ‰, where higher values indicate a higher dietary proportion of angling bait (Britton et al., 2022). These δ^13^C_corr_ values were highly correlated with TP (range 1.5 to 1.9), with fish of enriched δ^13^C_corr_ being of higher TP (linear regression: R^2^ = 0.75, F_1,8_ = 24.35, P < 0.01). We posit that fish with enriched δ^13^C_corr_ and of higher TP have higher angling recapture probabilities (due to their higher reliance on angling baits in the diet) and, accordingly, significantly higher scale cortisol content (due to multiple capture events in preceding months).

### 3.2 Capture vulnerability - scale cortisol relationships

Across the 12 barbel groups, differences in fork length between the groups of angled and electric-fished fish were minor (639 ± 34 vs 575 ± 49 mm), with this also the case in δ^13^C (−22.65 ± 1.39 vs -23.98 ± 1.32 ‰) and δ^15^N (12.16 ± 0.58 vs 12.83 ± 0.61 ‰). Scale cortisol concentrations per group ranged between 1.60 and 5.89 pg mg^-1^ (mean 3.17 ± 0.74 pg mg^-1^), with concentrations being substantially lower in groups that were captured by angling versus groups captured by electric fishing (2.68 ± 0.71 vs 4.14 ± 1.33 pg mg^-1^). This difference in scale cortisol concentration between angled and electric fished barbel was significant (GLM: Wald χ^2^ = 5.85, P = 0.02), where the effects of mean δ^15^N as a covariate in the model was significant (P = 0.03), but with all other covariates having non-significant effects (year of sampling: P = 0.06; mean length: P = 0.59; δ^13^C: P = 0.08; Fig. 1).

**Figure 1.**
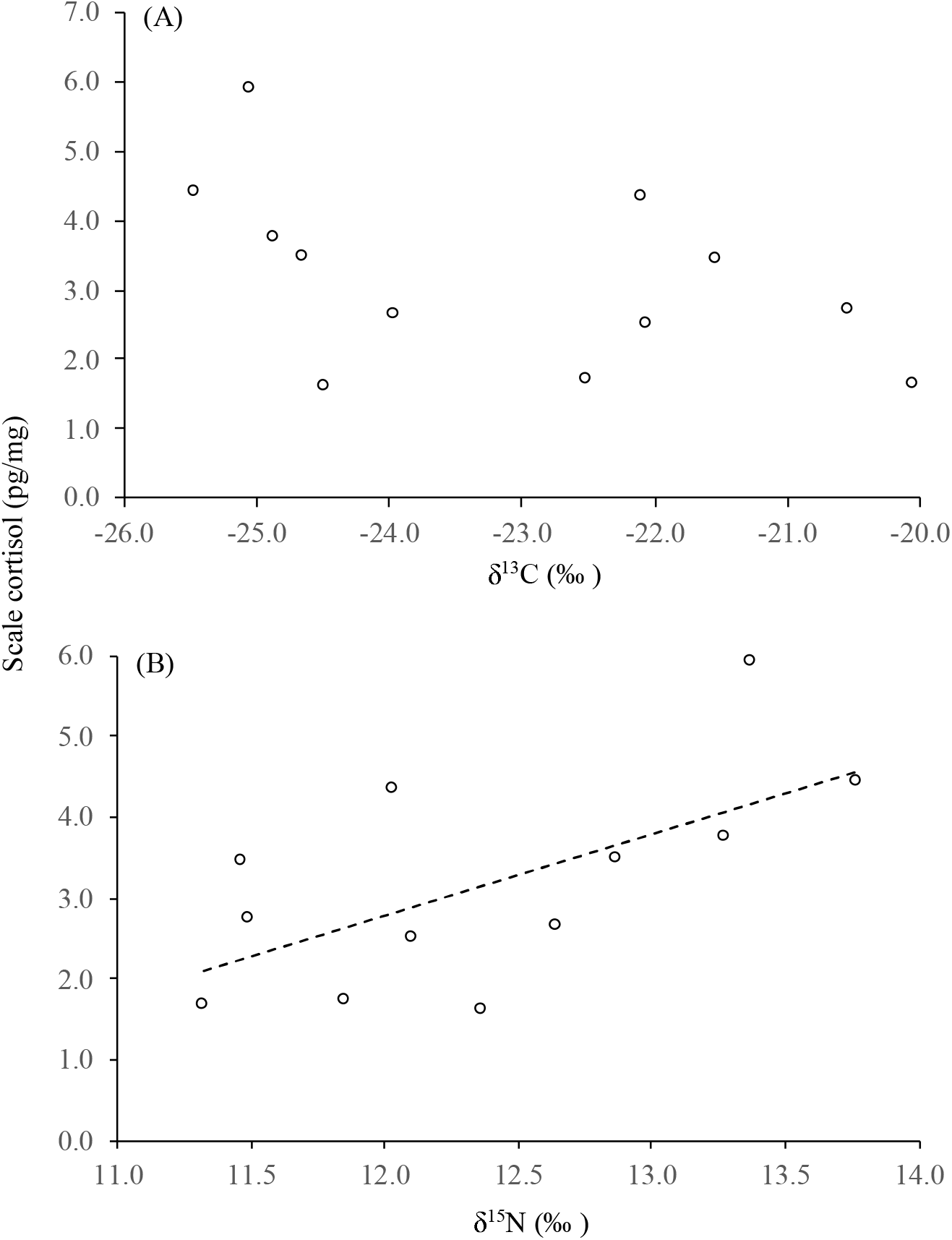
Relationship of (A) mean δ^13^C and (B) mean δ^15^N versus with scale cortisol concentration in pooled European barbel. In (B), the dashed line is the significant relationship between the variables according to linear regression (R^2^ = 0.40, F_1,10_ = 6.54, P = 0.03). In (A), the relationship between the variables was not significant (R^2^ = 0.23, F_1,10_ = 3.03, P = 0.11).

In the angled common carp, scale cortisol concentrations ranged between 1.26 and 5.40 pg mg^-1^ (mean 3.23 ± 1.26 pg mg^-1^). Linear regression indicated that as scale cortisol levels increased in individual carp, their values of both δ^13^C_corr_ and TP decreased significantly (δ^13^C_corr_: R^2^ = 0.57, F_1,8_ = 10.41, P = 0.01; trophic position: R^2^ = 0.55, F_1,8_ = 9.82, P = 0.01; Fig. 2). However, the relationship between fish length and scale cortisol concentration was not significant (R^2^ = 0.01, F_1,8_ = 0.09, P = 0.77).

**Figure 2.**
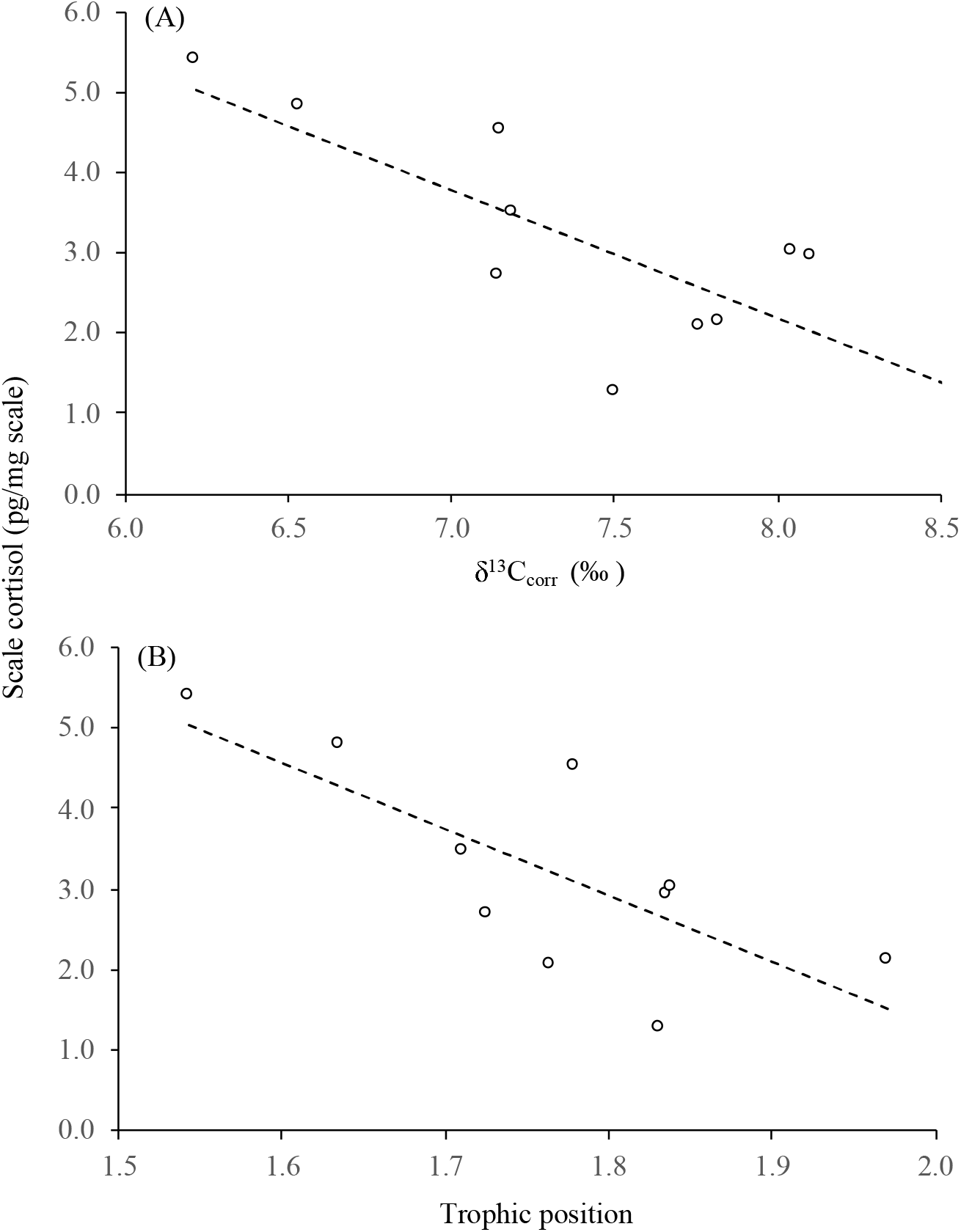
Relationship of (A) corrected carbon (δ^13^C_corr_) and (B) trophic position with scale cortisol concentration in common carp. Dashed line is the significant relationship between the variables according to linear regression (*cf*. Results).

## 4. Discussion

Scale cortisol content has been demonstrated as providing a reliable biomarker of chronic stress across a range of different fish species from both freshwater and marine environments (e.g. Carbajal et al., 2018, 2019a,b; Laberge et al., 2019; Lebigre et al., 2022l Vercauteren et al., 2022). Applying this biomarker here to two freshwater fishes exposed to catch-and-release angling, we initially developed proxies of angling capture vulnerability for the two study populations, where we assumed that fish with higher capture vulnerabilities would have had higher recapture rates in preceding months and, given individual angling capture events represent a short-term physiologically stressful event (Cooke et al., 2013), we hypothesised that these fishes would then have higher accumulations of cortisol in their scales. However, in both barbel and carp, significantly lower scale cortisol content was measured in the fishes that had been hypothesised as most vulnerable to angling capture. This result could have related to several factors, including fish captured regularly by angling have high stress resilience to repeated capture that is independent of other behaviours, fish with high capture vulnerability developing high hook avoidance behaviours following a capture event and/ or the proxies developed here on capture vulnerability did not adequately describe angling capture vulnerability in the two species.

Angling is recognised as a non-random method of fish capture, where angling induced selection results in population sub-groups with specific trait combinations being most vulnerable to capture. In many species, the most vulnerable trait combinations involve high activity and boldness, which generally align to high stress resilience within proactive-reactive coping styles (Castanheira et al., 2017; Vindas et al., 2017; Villegas-Rios et al., 2018). For example, a phenotypic syndrome in rainbow trout was evident whose physiological responses to an experimental stressor (including cortisol levels) was relatively low, with these fish then having a higher vulnerability to angling capture than other phenotypes (Koeck et al., 2019; Monk et al., 2021). The acoustic tracking study of the barbel completed in the lower River Severn basin in 2015 indicated that there was some inter-individual variability in their behaviours, with some individuals having relatively small home ranges (some < 1km, suggesting a low activity, bold phenotype), with others having home ranges that were substantially larger (> 12 km, suggesting a high activity, shy phenotype) (Gutmann Roberts et al., 2019). Moreover, the fish with smaller home ranges were primarily sampled by angling, whereas those with larger home ranges were captured by electric fishing, where electric fishing is much less selective in fish capture than angling (Vehanen et al., 2013; Radinger et al., 2019). Moreover, we then demonstrated here that there were considerable dietary differences between angled and electric fished barbel, with angled fish having substantially enriched δ^13^C versus electric fished barbel. Thus, we posited that barbel sampled by angling would have higher angling recapture rates than those electric fished, due to their smaller home ranges that increased their spatial encounters with anglers (Gutmann Roberts et al., 2019), and their higher dietary reliance on angling bait suggesting they would be more likely to encounter a baited hook while foraging. However, the barbel groups captured by electric fishing had significantly higher scale cortisol content than angled groups captured by angling, suggesting that if the electric fished barbel were a high activity phenotype, then this high activity was not associated with higher levels of stress resilience.

Although angling capture is recognised as a stressor that can have considerable short-term physiological effects (e.g. Cooke et al., 2013; Pinder et al., 2019), the chronic effects of angling are less apparent. Repeated capture in C&R angling was suggested as decreasing the growth of striped bass *Morone saxatilis* through the effects of released fish not resuming feeding for up to two days (Stockwell et al., 2002). In largemouth bass *Micropterus salmoides* and smallmouth bass *Micropterus dolomeiu*, Cooke et al. (2002) suggested that the repeated handling of fish during tournament angling (that involves multiple periods of handling and air exposure) is likely to impact on the biological fitness of individuals, with nest guarding males that are captured and released also being impacted through a reduced ability to nest-guard (due to reduced locomotory activity). Although knowledge of how multiple captures affect individual fish behaviour and physiology is more limited, evidence suggests that fishes which are captured can often then demonstrate ‘hook avoidance’ behaviours that reduces their subsequent capture vulnerability. For example, Raat (1985) demonstrated that after carp were hooked once, their vulnerability to subsequent capture was decreased, with experiments by Lovén Wallerius et al. (2020) indicating that in addition to direct experience of angling capture leading to hook avoidance, carp can also develop these behaviours through social learning alone. Although domesticated strains of carp are bolder and more vulnerable to angling capture than wild strains, the development of hook avoidance behaviours are similar across both strains (Kleforth et al., 2013). Thus, while our proxies for angling capture vulnerability were considered as likely to result in higher capture rates (and so higher scale cortisol content), it could be that following an initial capture event, these fishes developed strong hook avoidance behaviours that enabled them to continue foraging on angling baits while reducing their risk of ingesting a baited hook. Conversely, given that it is clear from other studies that a minority of individuals within populations are capable of being recaptured on multiple occasions (e.g. European catfish; Britton et al., 2007) then in these individuals, there might be trade-offs between the ease of resource and energy acquisition from feeding on readily available angling baits (that reduces their foraging times) versus the elevated risk of incurring repeated angling capture events, whose severity might reduce with capture frequency through habituation. However, this habituation to repeated angling capture is highly speculative in the absence of supporting evidence.

A limitation of this study was that in the population that had the most appropriate sample size for robust testing of chronic stress responses (barbel, n = 68), the method used to determine scale cortisol content required scale mass in excess of that able to be provided by individual fish. The pooling of individual fish into groups according to their capture method, sampling year and stable isotope data was considered as the most appropriate way of dealing with this issue, but it is acknowledged that the reduction from 68 individuals to 12 groups resulted in a relatively coarse analysis that could have resulted in the barbel phenotypes that were most vulnerable to angling capture and so having the highest levels of scale cortisol content being mixed with individuals of lower vulnerability. The relatively low sample size of carp and lack of replication across water bodies also limits our ability to transfer these finding to other fisheries and species. Notwithstanding, there was consistency in the results across both species whereby those fishes that we predicted would be most vulnerable to angling capture being the groups and individuals with the lowest scale cortisol content, suggesting angling high vulnerability correlates with high stress resilience. However, we suggest that if this biomarker is to be used subsequently, studies need to consider measuring capture rates more precisely in individual fish, with individual level material and data then used throughout the laboratory protocol and data analyses. In addition, the use of fish from waters where angling is not practised would provide samples that have not been exposed to this specific stressor.

In summary, the use of scale cortisol as a biomarker of chronic stress indicated that fish of high angling vulnerability had significantly lower scale cortisol content. In barbel, this was despite these fish being of a low activity phenotype that had been assumed to have lower stress resilience than the high activity phenotype (which had higher scale cortisol content). The reasons for these more vulnerable fish having lower scale cortisol remain unclear, but could relate to fish that are recaptured regularly being able to cope with the physiological demands of angling capture more easily than other fishes or these vulnerable fishes developing hook avoidance behaviours that minimises their recapture rates while enabling them to continue expressing other behaviours.

## Data sharing

Raw data used in the manuscript that are not already provided are available from the corresponding author on reasonable request.

## CRediT authorship contribution statement

JRB and DA conceptualised the study and provided scale samples, AC and MLB designed the laboratory protocols and completed all laboratory analyses, JRB analysed the data and wrote the manuscript, all authors revised the manuscript and all authors agree to its submission.

